# The prevalence of vitamin D deficiency between Saudis and non-Saudis in Al-Madinah Al-Munawarah a cross-sectional study

**DOI:** 10.1101/613729

**Authors:** Muhammed Hassan Nasr, Noordin Othman, Bassam Abdulrasol Hassan, Mahmathi Karoppannan, Noorizan Binti Abdulaziz, Mohammed Ahmed Alsarani, Mohammed Husain Eskembaji

**Affiliations:** Clinical Pharmacy Department, Faculty of Pharmacy, Taibah University, Al-Madinah Al-Munawarah, Saudi Arabia; Quality Use of Medicines in Umrah and Hajj Pilgrimage Research Group, Department of Clinical and Hospital Pharmacy, Faculty of Pharmacy, Taibah University, Al-Madinah Al-Munawarah, Saudi Arabia; Clinical Pharmacy Department, Faculty of Pharmacy, Universiti Teknologi MARA, Selangor, Malaysia; Clinical Pharmacy Department, Faculty of Pharmacy, Management and Science University; Laboratory Department, Medical Care Unit, Taibah University, Al-Madinah Al-Munawarah, Saudi Arabia

## Abstract

**Background:** Vitamin D, or the “sunshine” hormone became an attractable topic that recently captivates many researchers. The increased prevalence of vitamin D deficiency became an alarming health concern despite the accumulative evidence exploring its crucial role not only in bone metabolism, but also in a variety of pleiotropic functions throughout the various body organs. The aim of this study is to compare the prevalence that might influence vitamin D deficiency among Saudi and non-Saudi nationalities in Almadinah Almunawarh, Saudi Arabia, and to study the different factors that may have an influence in the difference of this prevalence like the marital status, occupation, smoking, sunlight exposure, education, and dietary habits.

**Methods:** The study was a cross sectional study done in the medical care unit in Taiba University Almadina Almunawarah in which, 65 healthy male individuals from different nationalities (Saudis and non-Saudis), aged 18 - 65 years were divided into 2 groups, 33 Saudis and 32 non-Saudis. A sociodemographic questionnaire was filled by the study participants and 25-OH vitamin D3 (25(OH)D3) concentrations were detected by electrochemiluminescence immunoassay.

**Results:** Results showed a Significant percentage of the participants in the Saudi group (n = 30, 91%) suffered from deficiency in vitamin D levels [25 (OH) D < 20 ng/ml] 12.57 ± 4.82 (mean ± SD), compared to only 47% (n = 15) in the non-Saudi group [21.56 ± 6.82 (mean ± SD)]. Vitamin D deficiency was found to be significantly higher in the Saudi group than the non-Saudi group with P = 0.001.

**Conclusion:** Results showed a significant increase in vitamin D deficiency in Saudi population than the non-Saudis P = 0.001. The occupation status was found to be the only factor positively correlated with vitamin D deficiency.

## INTRODUCTION

Vitamin D is one of the vitamins that are soluble in fat. It is also classified as a prohormone steroid, (1). Vitamin D has important functions in the endocrine, paracrine and autocrine systems, as so, it is regarded as the “sunshine” vitamin (2,3). Naturally, only two forms of vitamin D are available; vitamin D2, also known as ergocalciferol, and vitamin D3 (cholecalciferol). Photosynthesis of vitamin D in the skin by the induction of sunlight produces only cholecalciferol, while the dietary sources of vitamin D can provide the vitamin in the two forms (3).

Globally, Vitamin D deficiency is among the major public health issues (4,5). The deficiency in vitamin D in children leads to a disease called rickets in which bone tissue fails to mineralize properly, causing skeletal deformities and brittle bones (1,6–8). While, the deficiency in vitamin D in adults leads to weakness in the muscles and consequently, increasing the risk of falls and fractures. In adults, vitamin D deficiency causes another disease called osteomalacia, leading to the weakness of bones and exacerbate osteoporosis (1,6).

Numerous data have been documented on the link between ethnicity, colour of skin and vitamin D serum levels (9,10). In the United States, 90% or more of the non-Hispanic African American were found to have vitamin D levels around 15 ng/mL, the mean vitamin D level of Hispanics was 20 ng/mL, while that of the non-Hispanics white was 26 ng/mL, whereas those individuals residing the traditional counties in Central Africa were showed to have mean plasma 25(OH) D levels of about 46 ng/mL (1,11).

In the Middle East and North Africa (MENA), especially the Kingdom of Saudi Arabia (KSA), high prevalence of vitamin D deficiency has been reported despite the abundance of sunshine (12,13). In 2015, a cross-sectional national multistage surveyreported that the prevalence of vitamin D deficiency was 40.6% in male Saudis and 62.65% in female Saudis (14). Similar results were found in other, studies that were conducted in various places in KSA. Two studies that were conducted in Riyadh indicated that vitamin D deficiency among Saudi men was 87.8% (15), and;78.1% in Saudi females and 72.4% in Saudi males respectively (16). In Damam the prevalence of vitamin D deficiency was found to be greater than 65% (17). Furthermore, Dabbour and colleagues (2016a study) conducted in Makkah estimated that vitamin D levels were very low among the healthy Saudis population (5.26 ±2.59 ng/ml) (4).

To our knowledge, up to know no study has yet been conducted to compare the prevalence of vitamin D deficiency between Saudis and non-Saudis living in the same area. This will be the first study to document Vitamin D status in Al-Madinah Al-Munawwarah. Also, this paper will examine predictors of such deficiency.

## Methods

### Subjects and Study Design

This was a cross-sectional study conducted in Al-Madinah Al-Munawarah, KSA from October 2017 to May 2018. This period of eight months was chosen to reduce seasonality bias as June, July, August and September are the hottest summer months in KSA. The ethical approval was obtained before the beginning of the study; in October 2017, from Taibah University College of Dentistry Research Ethics Committee (TUCD-REC), which is the ethical review board in Taibah University.

Non-diabetic male Saudis and non-Saudis nationalities age between 18 to 65 years old attended Medical Unit, Taibah University were included in this study. Participants were excluded from the study if they consume vitamin D supplementations, diagnosed with renal, liver or cancer, thyroid or parathyroid diseases and receiving any drug interacting with vitamin D such steroids, orlistat, cholestyramine, phenytoin and phenobarbital. Based on the sample size calculation, a sample size of 65 encounters per study arm; 33 in group one (Saudis) and 32 in group two (non-Saudis) would detect the degree of difference with 80% power at α = 0.05.

Prior participants’ enrollment commenced, demographic data, a detailed history of diabetes (if present), socio-economic data, information about intake of vitamin D-containing diets, exposure period to sunlight per day were collected. Completion and return of written informed consent before participating in the study indicated voluntary agreement to participate in this study.

## Data Collection

### Anthropometric Data

The weight was measured in kilograms and rounded to the nearest 100 grams. Standard beam scale was used to measure the weight of participants. The participants weighed wearing light clothes and barefoot

A calibrated height board was attached to the scale to measure the height in centimeters. Weight in kilograms were divided by the square of height in meters to calculate the BMI. The body mass index (BMI) was used to evaluate obesity. Participants with BMI between 25 and 29.9 kg/m^2^ were considered overweight, participants with BMI between 30 and 39.9 kg/m^2^ were considered obese participants. While, when BMI found to be > 40 kg/m^2^ then they were considered morbidly obese. Determination of the waist circumference was done by measuring the broadest area between the edge of lower ribs and the iliac crest.

### Blood Sample Collection and Laboratory Analysis

Five millilitres of blood were collected by trained technicians. This was done under the supervision and guidance of the primary care physicians. Blood tubes were preserved in a cooler or refrigerator immediately. The time of preservation was not less than 30 minutes and did not exceed four hours before the technicians centrifuge them. The centrifugation process was done for about half an hour at 3000 RPM at 4°C. Following that, the technicians immediately separated the serum from the whole blood and freeze them at - 20°C. This was done at the biochemistry laboratory at the Medical Unit, Faculty of Medicine, Taibah University, Al-Madinah.

The 25-hydroxyvitamin D levels were measured by ECLIA assay by Cobas machine e 411. The level were considered as deficient (< 20 ng/ml), inadequate (20 − 30 ng/ml) and adequate (30 ng/ml) as recommended by the American Endocrine Society Clinical Practice Guidelines (2,18).

## Results

A total of sixty-five healthy participants were recruited in the study, subdivided into two groups, 33 Saudis and 32 non-Saudis. The sample size was estimated to provide 80% power to detect a difference in vitamin D deficiency between the 2 groups at a two-sided 0.05 significance level.

(25-hydroxyvitamin D) levels less than 20 ng/ml were considered as deficient, while levels between 20–30 ng/ml were considered insufficient, and levels greater than 30 ng/ml were considered adequate vitamin D.

**Table 1.**
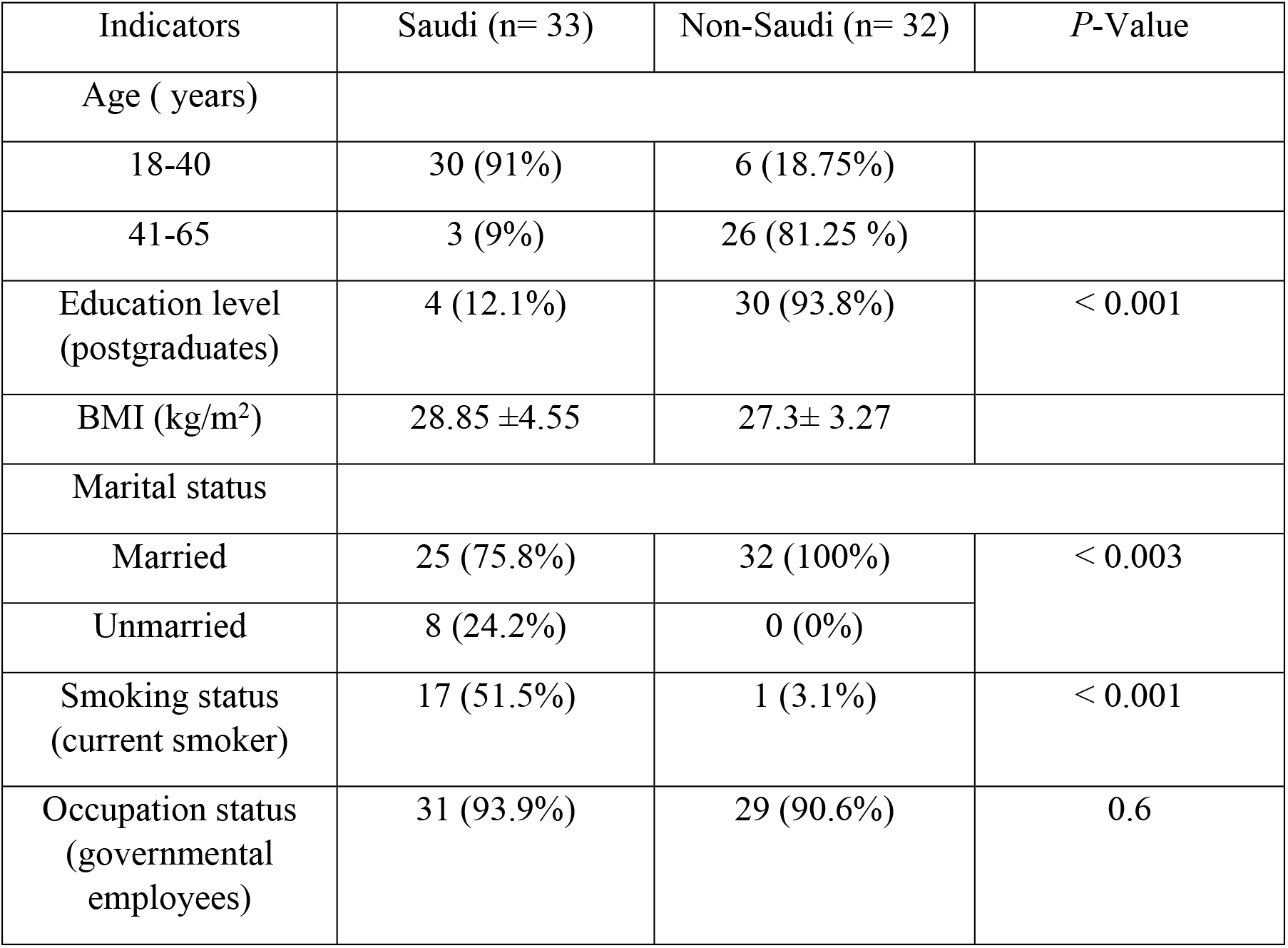
Baseline Demographic and clinical characteristics of participants.

The mean age of the Saudi group was 33.6 ± 7.2 years and the median and mode were found to be 32 years, while the mean age of the non-Saudi group was 46.8 ± 8.1 years, with median 45.5 and mode 43 years respectively.

### The marital status

All the non-Saudi group were married (100%) versus only 75.8% (n = 25) of the Saudi group were married. A significant difference was found between the two groups in their marital status.

### Smoking status

The results of the present study showed that high percentage of the Saudi participants were current and active smokers (n=17, 51.5%) compared to the non-Saudi participants (n=1, 3.1%). Statistical analysis showed that there is a significant variation in the smoking status between Saudi and non-Saudi participants (*p* < 0.001).

### Education status

Majority of the non-Saudi participants were post graduates, (n = 30, 93.8%) while only 12.1% (n = 3) in the Saudi group was postgraduates. Therefore, a significant variation was detected between the two groups in their education status (*p* < 0.001).

### The occupation status

The majority of both groups were governmental employees i.e. 93.9% (n = 31) Saudi and 90.6% (n = 29) non-Saudi, while only 6.2% (n = 2) of Saudi and non-Saudi participants were students. Besides that, 3.1% (n = 1) of the non-Saudi group were without a job. However, the occupation status did not differ significantly between the two groups (*p* = 0.6).

Majority of the non-Saudi participants were Egyptians (n = 19, 59.40%), followed by the Sudanese and Yemenis were 9.40% (n = 3) each, as shown in figure 1.

**Figure 1.**
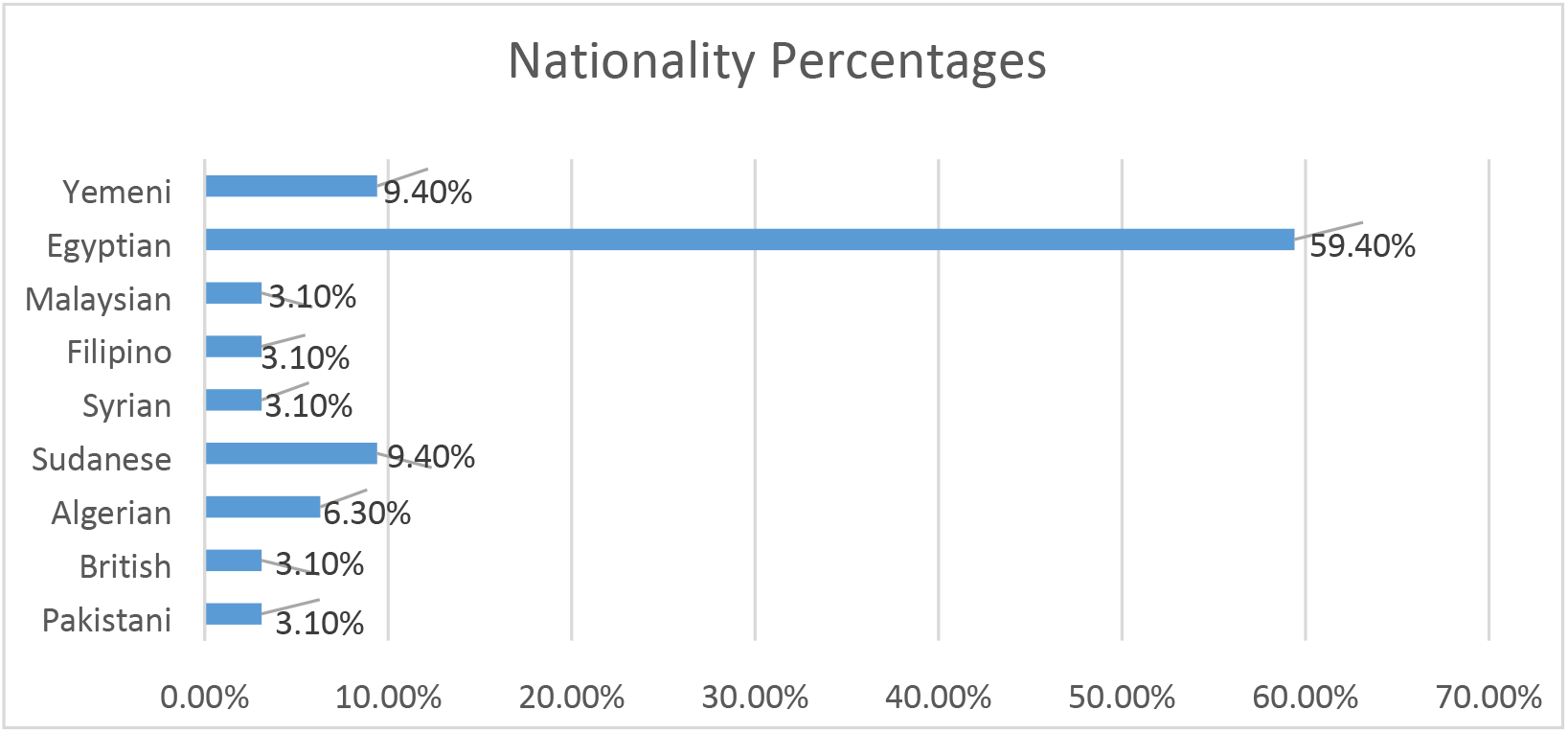
Nationalities of the non-Saudi participants (n=32)

**Figure 2.**
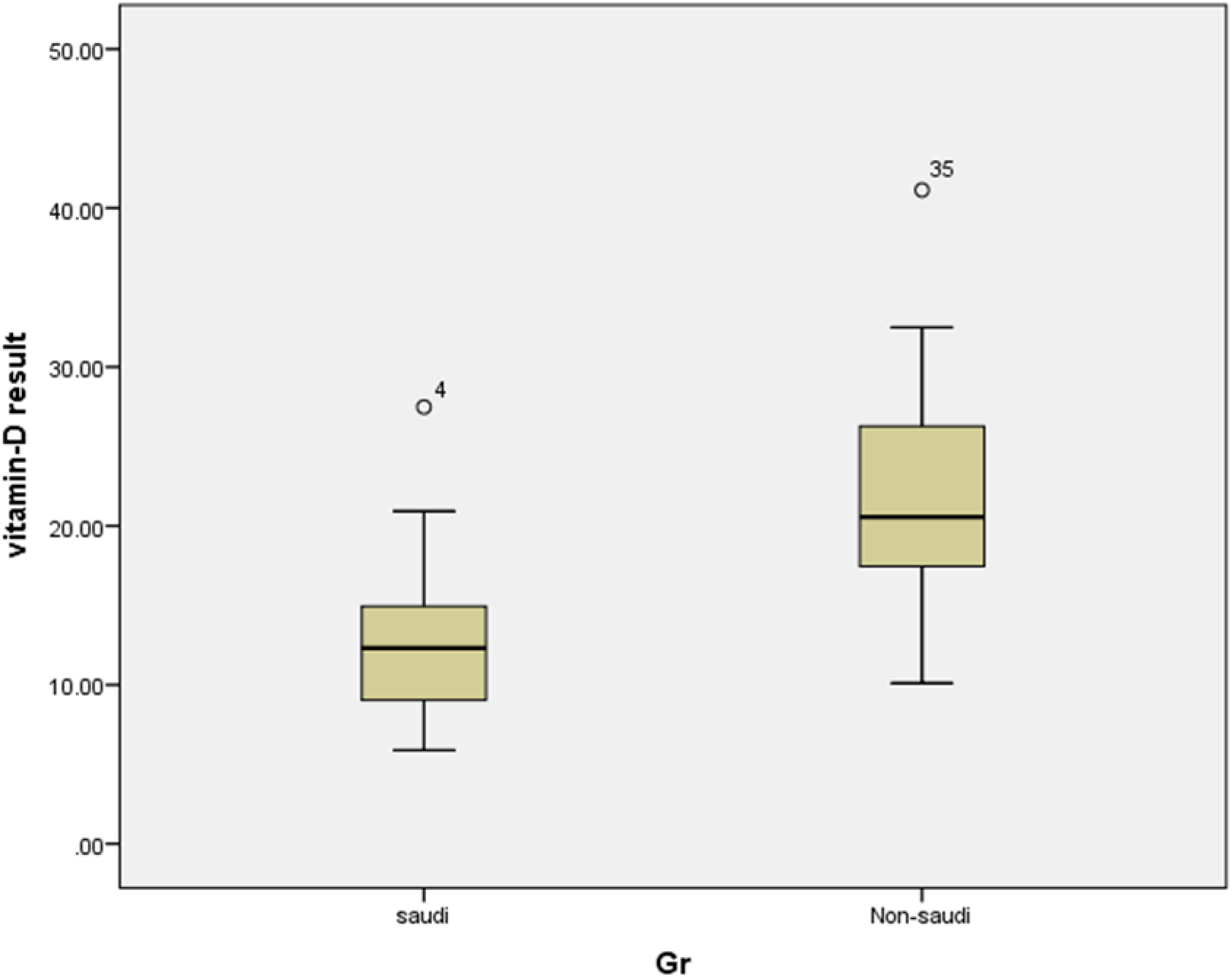
Box and whisker plots showing Vitamin D levels.

**Table 2.**
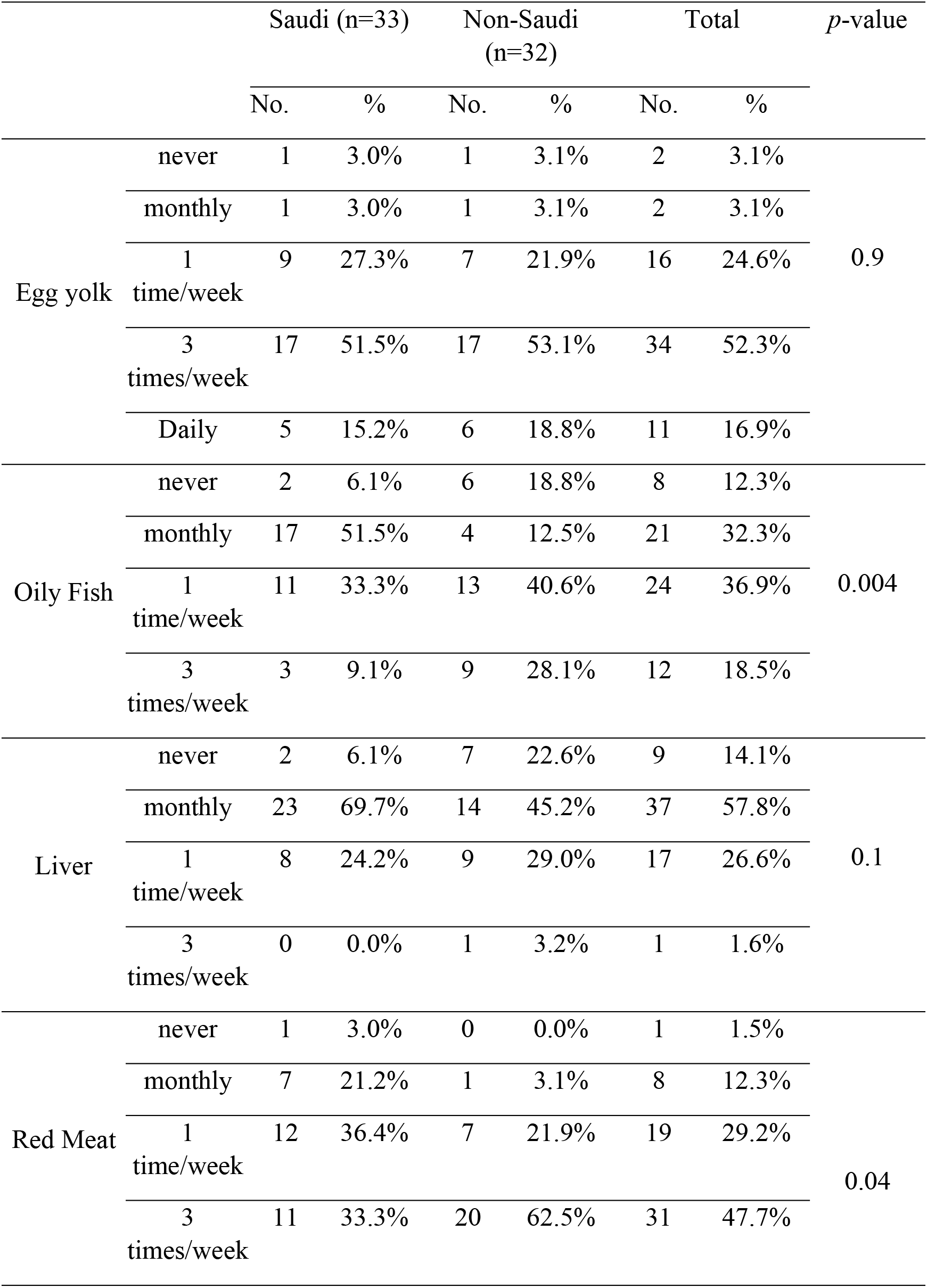
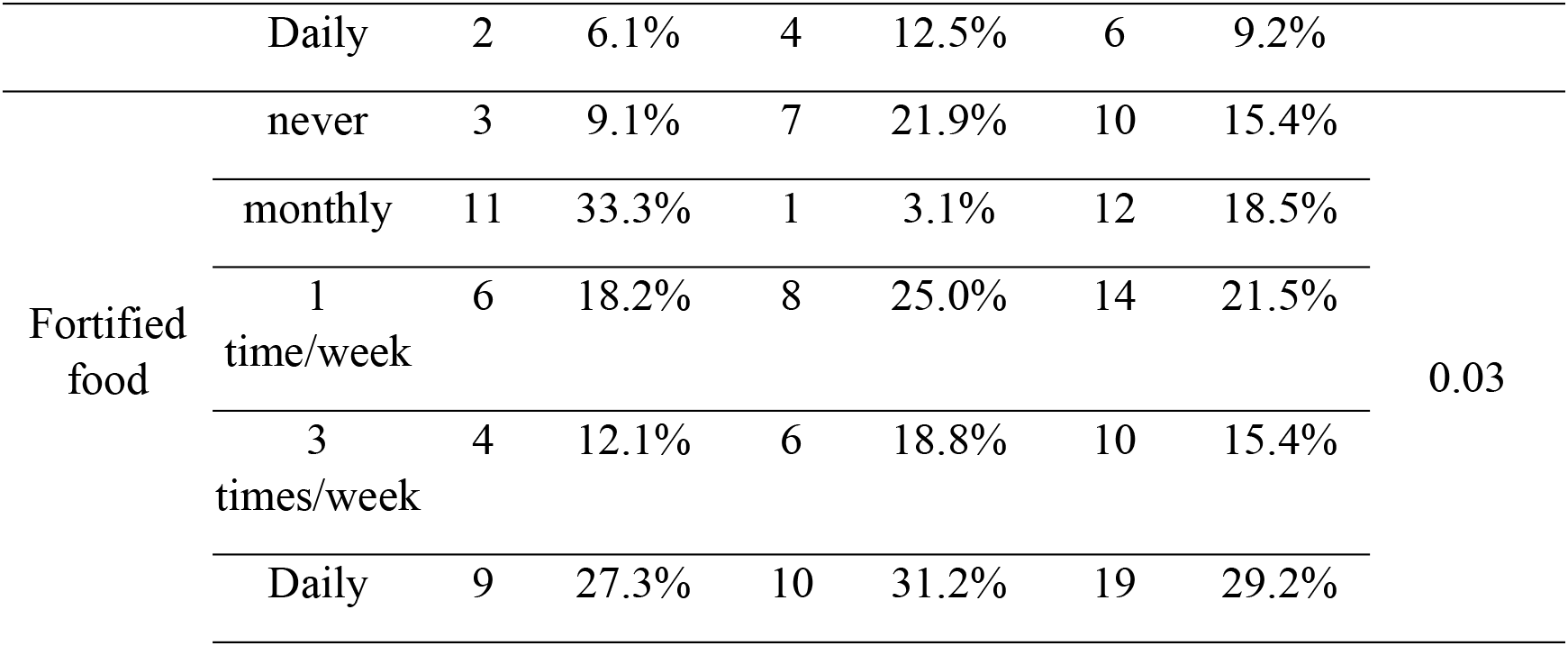
Dietary habits for the Saudi and non-Saudi participants.

Results of this present study showed that egg-yolk consumption in both groups were almost the same (51.5% versus 53.1%, 3 times/week, 3% versus 3.1% monthly, 15.2% versus 18.8% daily).

While, the statistical analysis of oily-fish consumption showed that non-Saudi population consumed more significantly than Saudi population did. There was not much difference between the two groups regarding liver intake, but the ingestion of both red meat and fortified food were significantly lower among the Saudi population compared with the non-Saudi.

**Table 3.**
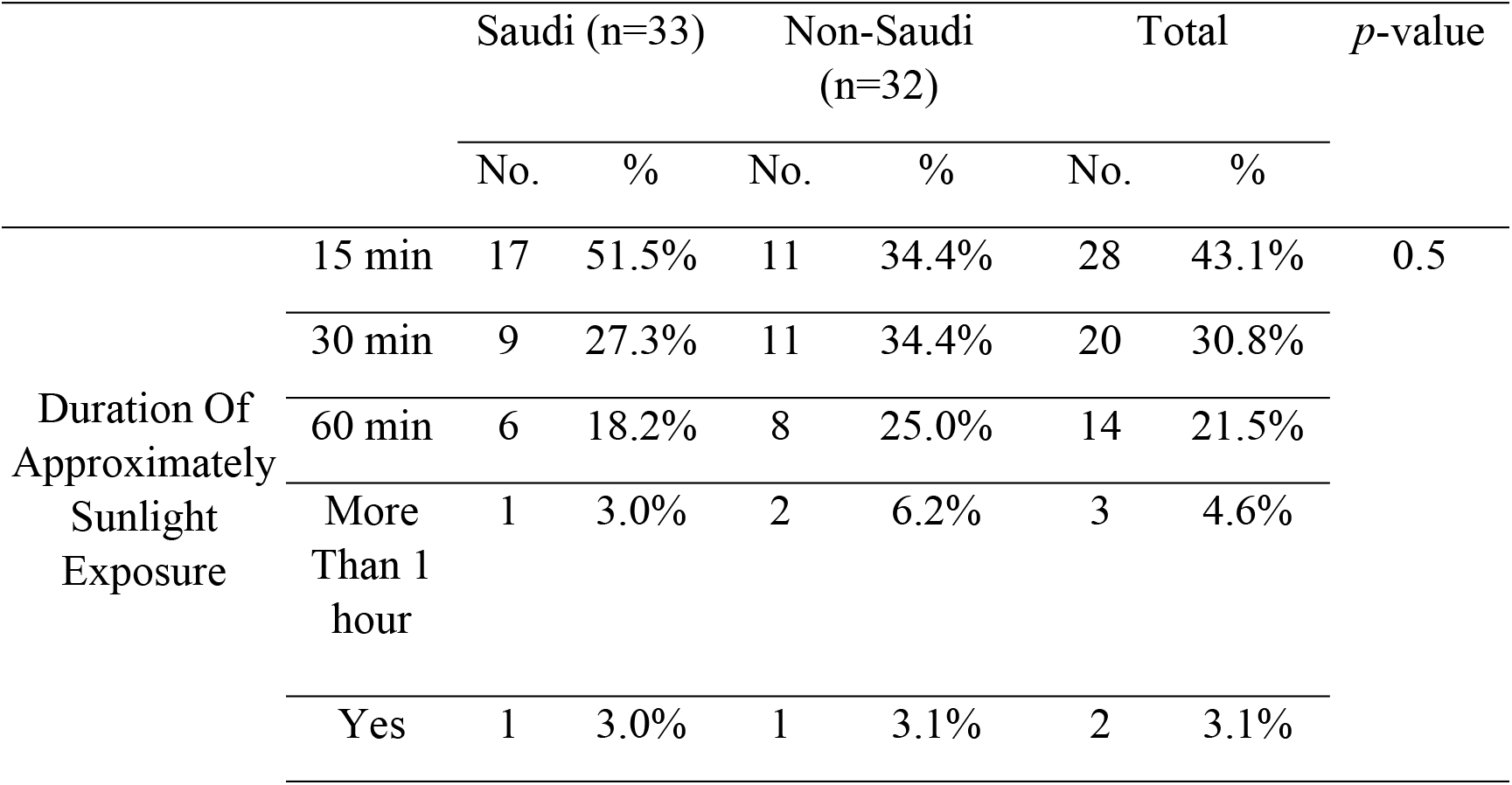
Duration and pattern of sun-light exposure.

In the Saudi nationality group, it was observed that 27.3% were exposed to sunlight 15 minutes daily compared to 34.4% in the non-Saudi group, while 18.2% of the Saudi nationality group were exposed to the sunlight 60 minutes daily versus 25% in the non-Saudi group.

Noteworthy, the positive exposure to sunlight in this study was defined as; the direct exposure (not indoors via windows) of at least some uncovered body parts, such as the arms and some parts of the legs, during the time period between 10 AM and 3 PM for not less than five minutes.

The occupation status was found to be positively correlated with vitamin D deficiency, while the consumption of liver was observed to be negatively correlated with vitamin D deficiency among the Saudi population, as revealed by logistic regression.

### Vitamin D levels

Significant percentage of the participants in the Saudi group (n=30, 91%) suffered from deficiency in vitamin D levels [25 (OH) D < 20 ng/ml] 12.57 ± 4.82 (mean ± SD), compared to only 47% (n=15) in the non-Saudi group [21.56 ± 6.82 (mean ± SD)].

Vitamin D deficiency was found to be significantly higher in the Saudi group than the non-Saudi group with *P* = 0.001

## Discussion

### Anthropometric Features and Socio-demographic data

#### Age of the participants

In this study, the ages of the Saudis were 33.63 ± 7.17 years (mean ± SD) with median and mode of 32 years. Meanwhile, the ages of the non-Saudis were 46.78 ± 8.1 years (mean ± SD), with median 45.5 and mode 43 years. Noticeably, the non-Saudis were in higher age group than the Saudi population in phase one of this study. That seemed reasonable because the majority of the non-Saudis (93.85%, n = 30) had postgraduate degrees; this process took a long time – around six to eight years after graduation. In addition, the main reason for the presence of the non-Saudis in KSA is for work, which they would not have been eligible for without at least three years of experience. All these justified why the non-Saudis were found to be in a higher age group than the Saudis.

On the contrary, the majority of the Saudi participants were without postgraduate degrees. All of them were local and working in their own country, which made it easier for them to be offered jobs in a short time just after graduation. In addition, most of the Saudi participants were administrative employees, for which postgraduate studies were not a must, so opportunities for governmental jobs were easy in their country. So, the ages of the Saudis were less than that of the non-Saudis.

This finding was consistent with that of Naugler et al. (2013) who observed that vitamin D deficiency was less prevalent in the older age group (19). However more than one study has demonstrated the high prevalence of vitamin D deficiency in different age groups among the Saudi population (20–22). The documented high prevalence of vitamin D deficiency among the Saudi population with wide age groups was mainly the result of the recent huge socioeconomic change that led to urbanization of lifestyles and westernization of dietary habits of most of the Saudi population.

#### BMI of participants

The BMI of the Saudis was 28.85 ± 4.55 (mean ± SD), while that of the non-Saudis was 27.3 ± 3.27 (mean ± SD). No statistical difference was detected between the two groups. The present study has shown that the Saudi population was characterized by higher BMI. Alqarni (2016) showed in a qualitative review an increasing pattern in the overweight and obesity prevalence in Saudi Arabia, which was expected to further increase in the future. Unbalanced diets – high consumption of carbohydrates in the form of rice, bread, pizza and dates with low vegetables and fruit intake –, in addition to the lack of physical exercise, contributed to the higher BMI and overweight status of the Saudi population (23–25).

In agreement with this finding, AlHumaidi, Adha, and Dewish in 2013 reported that obese Saudis with BMI 32.6 ± 6.7 (mean + SD) were found to have very low vitamin D levels (11.1 ± 5.9 ng/mL). Similarly, Al-Daghri et al. (2016) showed that overweight participants with BMI 28.0 ± 6.6 suffered from either vitamin D insufficiency (51.5%) or deficiency (29.9%) (26,27).

Also, the Saudi group had a greater deficiency in vitamin D than the non-Saudis. High BMI or obesity and vitamin D deficiency in this research was consistent with other researches that have evaluated the link between obesity and vitamin D status in different populations.

Bettencourt et al. (2018) performed a research in Portugal to assess the vitamin D status and its related factors. Their results showed that vitamin D levels were negatively correlated with obesity (i.e. BMI). This was mainly because of the sequestration of vitamin D in the body fat compartment, which could lead to the reduction of the bioavailability of vitamin D that was either photosynthesized through the skin by direct exposure to sunlight or from dietary sources (28). Other researchers have revealed the correlation of obesity with hypovitaminosis D with enhanced catabolism of the said vitamin in fat tissues and their conversion into inactive metabolites (29).

Kaddam et al. in 2017 conducted a cross-sectional study to assess the prevalence of the aforementioned condition and its associated factors among students and employees in three regions of KSA. The authors observed that obesity was associated with vitamin D deficiency (30). Tønnesen and colleagues in (2016) observed that obese participants had the highest relative risk for vitamin D deficiency in their study (31). Vimaleswaran et al (2014) showed that for every 1 kg/m^2^ higher in BMI, vitamin D deficiency was increased by 1.15% (32). Furthermore, Barchetta and colleagues in 2011 found that higher vitamin D deficiency was accompanied by increasing BMI and waist circumference (33). Chakhtoura and colleagues (2018) documented in their review on the prevalence of vitamin D deficiency in the Middle East and North Africa that higher BMI was one of the predictors of reduced serum vitamin D levels (13).

Therefore, it is important to consider much higher doses (2.5 times) of vitamin D supplementation in the obese vitamin D-deficient individuals due to their reduced bioavailability of vitamin D and/or the enhanced catabolism of vitamin D, as well as their conversion into inactive metabolites (1,18).

#### Marital status

The results of phase one in the present study have shown that all of the non-Saudi group and most of (75.8%, n = 25) the Saudi group were married (*P* = 0.003). As described before, the non-Saudi participants were of the higher age group than the Saudi participants. So, not surprisingly, the percentage of married non-Saudis was higher than that of the Saudis.

This finding could point towards the impact of marital status and social stability on the type of food, which might be healthier and containing a higher content of vitamin D. On the contrary, the unmarried individuals usually consume more fast and junk food that contained very small amounts of the vitamin.

#### Education status

The results have showed that the education status of the non-Saudi group was significantly higher than that of the Saudi group, where 93.85% (n = 30) of the non-Saudis had postgraduate degrees as compared to only 12.1% (n = 4) in the Saudi group (*P* = 0.001). This is because the administrative professions were restricted to the local residents (Saudi nationality) in KSA while the academic professions were mostly occupied by non-Saudis.

It is elicited in this present study that participants with higher education levels had higher vitamin D levels. The same observation was seen in a number of previous studies. In the Emirates, Bani-Issa and colleagues performed a quantitative cross-sectional study in 2017 to assess vitamin D deficiency and the factors associated with this condition. The authors have concluded that the less-educated, employed Emirati participants had a significantly higher percentage of vitamin D deficiency than the higher-educated participants (12).

In Alberta, Canada, an ecological study has been conducted by Naugler et al. (2013) to evaluate the association between vitamin D deficiency and a number of socio-demographic factors. The authors have observed that vitamin D level was associated with education status, and they attributed their finding to the increased consumption of vitamin D supplements by the higher-educated individuals (19).

Moreover, Ardawi and colleagues in 2012 studied the high prevalence of vitamin D deficiency among the Saudi males. The authors attributed the significant deficiency in vitamin D among Saudis to a number of factors, among which was the lack of education (34). Similarly, a Finnish study performed by Jääskeläinen et al. (2012), has documented the association between vitamin D levels with the education status (35). Furthermore, in China, Song et al. (2013), who studied the prevalence of vitamin D deficiency in pregnant women, have observed that the long duration of sunlight exposure, and the subsequently alleviation of vitamin D deficiency was in favour of the higher-educated women (36).

Notably, high education level rendered the individuals to be more knowledgeable and have greater awareness of at least the basic role of vitamin D in body-functioning and the negative impact of its deficiency.

#### Smoking status

There was a significant variation in the smoking status between the two groups with *P* = 0.001, whereby 17 out of 33 (51.5%) from the Saudi group were current smokers, while only 1 out of 32 (3.1%) from the non-Saudi was a current smoker.

According to Al-Nozha and colleagues, smoking rates were high among Saudis who were living in urban areas (Al-Nozha et al., 2009). This was consistent with the finding of this present study, as Al-Madinah is one of the urban cities in KSA. (37).

Smoking status of the Saudi population could help explain the deficiency in vitamin D levels detected in this population. Chakhtoura and colleagues, in their systematic review in 2018, have reviewed trials that studied the vitamin D deficiency in the MENA region. They indicated that smoking status was one of the possible consistent predictors of vitamin D deficiency (13).

In addition, several studies that have been conducted in different countries and used different methodologies indicated similar findings regarding the link between smoking status and vitamin D levels (38–40).

Hoteit et al. (2014) who studied the prevalence of vitamin D deficiency in 9147 Lebanese subjects reported smoking status was a robust predictor of vitamin D deficiency (38). Jiang et al. (2016), via a cross-sectional study that was performed in China, observed that the current smokers were associated with more severe vitamin D deficiency than non-smokers (39). In addition, Tønnesen et al. (2016) considered smoking as one of the modifiable factors for vitamin D deficiency as they showed that smoker participants were associated with a higher relative risk of vitamin D deficiency than non-smokers (31).

Furthermore, in 2014, Mulligan and colleagues have assessed the impact of current smoking on vitamin D-serum levels. Their results showed that smoking caused a significant reduction in vitamin D serum levels (40). Ardawi and colleagues (2012) reported a high prevalence of vitamin D deficiency (87.8%) among Saudis living in Jeddah. The authors have shown that smoking was one of the attributable factors and was associated with the low vitamin D status found in the Saudi population (34). Notably, Brot and colleagues (1999) have conducted a cross-sectional study to investigate the effects of smoking on the levels of vitamin D and calcium metabolism. The authors have observed that smoking adversely affected calcium and vitamin D metabolism, mainly by depressing vitamin D-parathyroid hormone pathway (41).

Not surprisingly, that the results of all these studies were in agreement with this present finding regarding the link between smoking and low vitamin D levels, as it has been documented that smoking negatively affected various body functions, harmed organs, and had a lot of general adverse consequences throughout all the body (42).

#### Occupation status

There was no significant difference in the occupation status between the two groups (*P* = 0.60), as 93.9% (n = 31) from the Saudi group and 90.6% (n = 29) from the non-Saudi group were governmental employees. The participants in the two groups were employees in Taibah Univeristy, which is a governmental university in Al-Madinah Al-Munawarah. Being office-bound employees had a negative impact on their lifestyle, rendering them to have more sedentary lifestyles and decrease their exposure to sunlight – which is the major source for the photosynthesis of vitamin D. Tønnesen et al. (2016) have documented that the more the time spent in physical exercise, the more reduction in the relative risk of vitamin D deficiency. The authors have also added that fast food consumption was associated with a high relative risk of vitamin D deficiency (31). Furthermore, Dabbour and colleagues performed a research in Makkah region in Saudi Arabia in 2016 to evaluate the association of vitamin D deficiency with T2DM. The authors have observed that vitamin D deficiency – which is highly prevalent in KSA – was associated with the lack of physical activity and decreased exposure to sunlight. They further added that physical activity could aid in the maintenance of vitamin D levels, hence concluding that duration of physical activity was positively correlated with serum vitamin D levels (4).

### Dietary habits of participants

Detection of dietary habits of the Saudi population could be considered as a very important point, because there is still scarcity in the information on vitamin D status and its association with the Saudi population’s dietary habits. Such information can help ministries in charge (e.g. ministry of health, ministry of education, and ministry of media) guide the Saudi population to overcome this medical problem (i.e. hypovitaminosis D), either by guiding the general population to increase exposure to sunlight or by focusing on the fortification of food products (43). The answers to these two points have been mentioned by this present study. Such valuable information could significantly help develop a guideline for the Saudi population.

The dietary habits between the two groups were found to be as follows:

**Oily fish**: the consumption of oily-fish was significantly higher in the non-Saudi group than in the Saudi group (*P* = 0.004).

Salmon reportedly contains 988 IU of vitamin D per 100 grams; this amount accounts for about 247% of the Reference Daily Intake (RDI) of vitamin D. Herring and sardines: the fresh Atlantic herring contains around 1628 IU of vitamin D per 100-grams; this amount accounts for around 4 times the RDI. Pickled herring contains about 680 IU/100 g that accounts for 170% of RDI. Sardines provide 272 IU per serving. Mackerel provides 360 IU/serving. In addition, cod liver oil is a good source of our concerned vitamin as it contains around 450 IU/teaspoon. Furthermore, canned tuna provides up to 236 IU/100 g (44).

Oily fish contains omega-3 fatty acids which have a well-known protective effect against dyslipidemia, hypertension, stroke, and cardiovascular diseases, in addition to the fair contents of protein, vitamins, and other essential nutrients for good health (Burger et al., 2014). The consumption of oily fish by the Saudi community remained poor. This could have been due to cultural and traditional reasons as the main food types in Saudi Arabia are mainly rich in carbohydrates and fibers like rice, potatoes, bread, and dates. In addition, of the diet types and habits have increasingly westernized in the last few years due to the introduction of multi-national restaurants in the Saudi market. Furthermore, the lack of health and nutritional education played a crucial role in the dietary habits of the Saudi population. These observations were consistent with those of Burger et al. (2014) who have studied the behaviour and rates of fish consumption among the natives and non-natives living in KSA. The authors have shown that the percentage of the male Saudis who consumed fish was 3.7% and that of the male non-Saudis was 6.6% (45). Kaddam et al. in 2017 conducted a cross-sectional study to assess the occurrence of hypovitaminosis D among students and employees in three regions of KSA and its causative factors. The authors observed that lack of omega-3 in the diet of the students was associated with vitamin D deficiency (30).

**Red meat**: the non-Saudi group consumed red meat significantly more than the Saudi group (*P* = 0.04), as 62.5% (n = 20) of the non-Saudis consumed red meat three times weekly, and 12.5% (n = 4) ingested red meat on a daily basis, as compared to 33.3% (n = 11) and 6.1% (n = 2) of the Saudi group who consumed red meat in the same pattern.

A study has documented the levels of vitamin D_3_ in types of meat, which were found to be 9.0 μg/kg for beef, 1.0−23.0 μg/kg for pork, 1.0−61.0 μg/kg for lamb, 0.0−50.0 μg/kg for veal, 0.0−14.0 μg/kg for poultry, and 0.0−23.0 μg/kg for various meat product (46).

As mentioned before, the inherited traditional dietary habits in the Saudi community led them to prefer foods that were rich in carbohydrates and fibre, and low in fats and cholesterol. Moreover, the socioeconomic conditions, which have changed in the last decade in Saudi Arabia, caused the migration of significantly large proportions of the Saudi population from the rural areas to settle in the urban and large cities. It has been forecasted by the Ministry of Municipal and Rural Affairs (MMRA) that by the year 2025, 88% of KSA residents will settle in the urban areas (47). Hence, the modification and urbanization of their lifestyles have led to the alarming increase in the consumption of fast and junk foods such as pizzas, pastas, and sandwiches. This explained the Saudis’ low consumption of red meat.

This present study’s finding was consistent with a nationally representative survey done in 2016 by Moradi-Lakeh and colleagues to evaluate the dietary habits in KSA. The authors have discovered that the type of food least consumed by the Saudi population was red meat; 4.8 ± 0.2 g (mean ± SE). They added that only a small proportion of the Saudis met the dietary recommendations (48).

**Fortified food**: Although, fortified food may be considered as the ideal and the easiest alternative to sunlight exposure in the Saudi lifestyle to guard against vitamin D deficiency, yet the consumption of fortified food was significantly higher in the non-Saudi group than in the Saudi group (*P* = 0.03).

The known sources of fortified food are cow’s milk which usually contains around 130 IU vitamin D/cup (237 ml), soy milk which can contain up to 119 IU vitamin D/cup, orange juice which can also be fortified with vitamin D (usually 142 IU/cup), and cereal and oatmeal which can be fortified with about 154 IU vitamin D per serving (49). This finding is consistent with two recent studies which were conducted in Saudi Arabia. Al-Daghri (2015) has studied the correlation between vitamin D levels and the consumption of dairy and fortified products in Saudi Arabia. The author has observed the poor consumption of fortified and dairy products in the overall Saudi population, apart from showed that the vitamin D status was significantly associated with the consumption of dairy and fortified products (20). In addition, in 2012, Ardawi and colleagues considered poor consumption of dietary and fortified products as one of the contributory factors to vitamin D deficiency in the Saudi population (34).

**Egg-yolk**: in contrast, there was no significant difference between the two groups in the consumption of egg-yolk (*P* = 0.9). Only a small proportion of the two groups ate egg-yolk as per the dietary recommendations. This could be justified by the dramatic socioeconomic change and hence, urbanization of lifestyle and westernization of food habits which pointed towards the large consumption of fast-food. Egg-yolk in general is rich in fats, vitamins, and minerals. Egg-yolk contains around 39 IU of vitamin D, while egg-yolk which comes from chickens that spent more time under the sun produced 3 to 4 times higher vitamin D. Chickens fed with vitamin D-enriched feed produced egg-yolks with around 6000 IU vitamin D (49).

**Liver consumption** did not vary significantly between the two groups (*P* = 0.1). Even though liver is considered as one of the rich animal sources of vitamin D that reaches as high as 140 μg/kg, the consumption of liver was quite poor in both groups (46).

**The duration of sunlight exposure** did not significantly vary between the two groups (*P* = 0.5). As observed, both groups were poorly exposed to sunlight (18.2% from the Saudi group and 25% from the non-Saudi group were exposed to sunlight for one hour daily).

It was expected that 80% of the daily requirement of vitamin D in the majority of individuals would be generated through sunlight exposure, as the major source of vitamin D was photosynthesis. However, this was not the reality nowadays, presumably due to the fear of erythema and some types of skin cancer, or that the increasing use of technology in the 21^th^ century kept most of the people spending most of their time indoors (1). In Saudi Arabia, apart from the abovementioned possible causes of the avoidance of exposure to sunlight, this could be attributed to the extreme hot temperature in KSA due to its geographical location, abundant use of cars, and the presence of more highways.

Consistently with our results, Ardawi and colleagues in 2012 assessed the vitamin D deficiency prevalence among healthy Saudi males. They have observed the high and significant prevalence of vitamin D deficiency in this population was because of a number of factors, among which included poor sunlight exposure (34). Almehmadi et al. (2016) studied the vitamin D levels among children in Jeddah, KSA. Their result – poor sunlight exposure in Saudi Arabia – was consistent with the finding of this present study (50). The discrepancy between both studies was in the study population, where this present research focused on adults and not children.

### Vitamin D levels among Saudi and non-Saudi populations

In phase one, 30 out of 33 participants in the Saudi group (91%) were vitamin D-deficient (calcidiol levels < 20 ng/ml) as compared to 15 out of 32 participants (47%) in the non-Saudi population. Subsequently, vitamin D deficiency was found to be significantly higher in the Saudis (12.57 ± 4.82) (mean ± SD) than the non-Saudis (21.56 ± 6.82) (mean ± SD), with *P* = 0.001.

In terms of the socio-demographic data, occupation status was positively correlated with the vitamin D deficiency in the Saudi group and was shown to be the only predictor of vitamin D deficiency, as revealed by logistic regression and expressed by exponential beta. This finding could be explained by the fact that the majority of the Saudi population in this present study were governmental employees, which rendered them more office-bound, and consequently reducing their duration and the time of sunlight exposure (51). Among the dietary habits, the logistic regression analysis revealed that the consumption of liver was negatively correlated with vitamin D deficiency and was considered as a predictor of vitamin D deficiency.

Previously, there was a controversy regarding the exact level of vitamin D which could be considered as deficiency or even insufficiency. In the renal and extra-renal tissues, the enzyme 1α-hydroxylase (CYP27B1) targets its substrate calcidiol and converts it into the active form, calcitriol. It has been documented that not more than 50% of maximal 25- (OH) D-1 α -hydroxylase activity (Km) was reached when 25-(OH) D level reached 40 ng/mL (100 nmol/L), which in turn relied on having adequate amounts of vitamin D (18). On the kinetics of 25-(OH) D-1 α -hydroxylase, the most recent studies were consistent with the USA Endocrine Society’s recommendations that the targeted level for optimal vitamin D effects was 30–50 ng/mL (75–125 nmol/L), preferably achieving the level of 40–60 ng/mL (100–150 nmol/L). Meanwhile, vitamin D deficiency was denoted by serum vitamin D levels of < 20 ng/mL (< 50 nmol/L), and insufficient or suboptimal status 20–30 ng/mL (50–75 nmol/L (18,52,53).

Notably, 13 out of 33 Saudi non-diabetics were overweight (BMI > 25), and 5 out of 33 were obese (BMI > 30). So, it was observed from the finding of present study that both conditions – overweight and vitamin D deficiency – were higher in the Saudi population. Furthermore, it is well known that T2DM became an increasingly widespread chronic disease in the Saudi community (as mentioned in the Introduction Chapter). Thus, overweight or obesity and vitamin D deficiency may be considered indicators or predictors for the development of T2DM among the Saudi population. Likewise, Bani-Issa and colleagues (2017) have observed obesity and T2DM to be bivariate correlators (which were independent of each other) with vitamin D deficiency in Emiratis (12). In addition, Sadiya et al. (2014) documented in their cross-sectional study which was performed in UAE the coexistence of these three cofactors – obesity, vitamin D deficiency and T2DM (54). Therefore, the present study recommends early screening and detection of T2DM in the presence of obesity and vitamin D deficiency, with rapid and aggressive corrections of these two predisposing factors.

As long as vitamin D deficiency is highly prevalent in the Saudi population relative to the non-Saudis, there might be some genetic factors playing a role in this scenario. Al Kadi and Sonbol studied the high prevalence of vitamin D deficiency among Saudi population in 2015. They have stated that the increased rate of consanguineous marriages in the Saudi population have led to some types of genetic mutations, such as calcium-sensing receptor gene A986S polymorphisms (55). In addition, Sonbol and colleagues have reported the association between low levels of vitamin D and the “S” of the calcium-sensing receptor gene (56).

Furthermore, in 2016, Nayak and Ramnanansingh in their study which investigated the deficiency of vitamin D among the Trinidadian population have attributed this deficiency to VDR genetic polymorphism. This polymorphism might lead to the dimensional conformation of the vitamin D receptors which showed variations in their affinities towards vitamin D (57). This hypothesis gave rise to the need for other studies to confirm this hypothesis. A possible explanation for the rampant hypovitaminosis D among Saudis could have been due to the dark skin, which could reduce the Ultraviolet B (UVB) rays’ penetration into the skin, thus minimizing the synthesis of vitamin D (58).

However, all the various factors that have been discussed previously in this study or in other studies (poor consumption of vitamin D-containing food, skin pigmentation, sunlight exposure, obesity, smoking, interacting drugs, etc.) which are implicated in the incidence of vitamin D deficiency could lead to a decrease in the synthesis and bioavailability, and elevations in catabolism and urinary excretion of vitamin D (1,59).

In addition, the cultural clothes used by the male Saudi population which covered the whole body (even their heads and necks) reduced their body surface exposure to sunlight. Consequently, their cutaneous vitamin D synthesis was decreased.

The findings explored in this current study were consistent with one systematic review and six studies conducted in different regions in Saudi Arabia. However, to our best knowledge until nowour study was the first to compare the prevalence of vitamin D deficiency between the Saudi population and participants from other nationalities living in the same environment. In addition, it was the first to assess the prevalence of vitamin D deficiency in Al-Madinah Al-Munawarah region.

The systematic review and a meta-analysis was performed by Al-Dagrhi and published in 2018. The author has reviewed the literature for the most recent epidemiological clinical trials that studied the prevalence of hypovitaminosis D in the Saudi population from 2011 till 2016. He included 13 local trials (performed in KSA), in which 24 399 healthy Saudi participants were included. This systematic review-cum-meta-analysis has shown that 81.0% of the Saudi population were deficient in this micronutrient, and that the deficiency in vitamin D in the Saudi population was accompanied with several diseases, among which were asthma, cancer, cardiac diseases, chronic kidney disease, obesity, metabolic syndrome, type 1 diabetes, type 2 diabetes, and skeletal effects. The author has concluded in line with our results in which there were widespread hypovitaminosis D that was associated with insulin resistance and related comorbidities in the Saudi populations (43).

Mogahed (2018) has conducted a study in Riyadh city among 100 Saudi patients to detect their vitamin D status. The results showed that 80% of the participants suffered from vitamin D deficiency (< 20 ng/ml). The author has concluded that prevalence of vitamin D deficiency in the Saudi population was very high (60).

Dabbour et al. performed a case-control study on 200 participants in Makkah Region in Saudi Arabia in 2016 to evaluate the levels of vitamin D and to assess the associated factors in healthy and diabetic Saudi participants. This study has concluded that there was a very significant deficiency in vitamin D among the healthy and diabetic Saudi populations (4).

However, this present trial differed from Dabbour et al. (2016) and Mogahed (2018) in the checking and comparison of vitamin D levels among Saudi and non-Saudi population, and in the evaluation of the dietary habits of the participants in an attempt to assess their correlations with vitamin D deficiency.

A retrospective study has been conducted by Alfawaz et al. in Riyadh city to determine the prevalence of vitamin D deficiency and its association with cardiovascular disease and other factors in the Saudi population. Their results have showed that the overall prevalence of vitamin D deficiency in their study was 78.1% in Saudi females and 72.4% in Saudi males. Moreover, the prevalence of vitamin D deficiency was associated with age, and obesity, and positively correlated with calcium, albumin, and phosphorus. It was negatively correlated with alkaline phosphatase and PTH levels (16). This present study showed a higher prevalence of vitamin D deficiency among the Saudi population in Al-Madinah Al-Munawarah (91%) than what was reported by Alfawaz et al. (2014) in Riyadh (72.4.1% in males). Furthermore, this present study has not only detected the prevalence of vitamin D deficiency among the Saudi population, but also compared the high prevalence with those of the non-Saudi population living in the same environment via a prospective study design.

A multi-centered case-control study has been done by Alhumaidi and colleagues in the Southern Region of Saudi Arabia (mainly Khamis Mushyt and Abha) in 2013. Their aim was to assess the vitamin D status among the Saudi population. They have included 172 diabetic Saudi patients and 173 non-diabetic Saudi participants in their trial. They have showed a very significant vitamin D deficiency in the healthy and diabetic Saudi population (98.5%). The authors have suggested the need for larger studies for better detection of vitamin D deficiency in the Saudi population as a whole, especially among the diabetic patients (26). The lifestyle, socio-demographic data and dietary patterns of their study population have not been studied by Alhumaidi and colleagues, in contrast to this present study. In addition, participants with thyroid diseases were included in their study; 23.5% were found to have euthyroid multinodular goiter and 33.8% were with hypothyroidism. These could have influenced their results (61). Furthermore, 17.4% of the non-diabetic Saudi participants were lost to follow-up. In the contrary, this present study has excluded participants with any thyroid and parathyroid disease and the compliance with the study protocol in this particular phase was 100%.

Also, a study has been done in Saudi Arabia in 2012 by Ardawi et al. This study design was cross-sectional and was aimed to assess the prevalence of deficiency in vitamin D in Saudi men in Jeddah. The authors have concluded that the deficiency in vitamin D was highly prevalent (87.8%) among Saudi men living in Jeddah (62). Although this present study was similar to that of Ardawi et al. (2012) in detecting the prevalence of vitamin D deficiency in only males and both have reported the high prevalence of vitamin D deficiency in Saudi men. Ardawi and colleagues neither compared this high prevalence in Saudi men with other nationalities residing in the same environment, nor evaluated the association of socio-demographic factors, sunlight exposure, and dietary habits with the high prevalence of vitamin D deficiency in the Saudi and non-Saudi population.

In Norway, a study has been performed in 2013 to compare the serum vitamin D levels among different populations living in Norway. The results of this study have revealed that immigrants from South Asia, the Middle East, and Africa were found to be more vitamin D-deficient (25-(OH) D < 20 ng/mL) than East Asians (63). Notably, the authors have recommended the routine measurement of vitamin D levels and early detection of vitamin D deficiency in these populations (Middle East and Africa).

## Conclusion

Vitamin D deficiency was significantly higher in the Saudi population than the non-Saudis population. The occupation status was found to be the only factor positively correlated with vitamin D deficiency.

Early screening for vitamin D serum-level is recommended for the Saudi population and rapid correction of vitamin D deficiency with vitamin D supplements should be considered.

## Limitations

Multi-centred study might have provided more generalizable results.

## Acknowledgments

We would like to thank Dr Sherif Monir for his advice on statistical analysis, Dr AbdelMannan Aman for the assessment of participants invited to this study all the laboratory staff for their help with the data collection.

